# An Osteocalcin-deficient mouse strain without endocrine abnormalities

**DOI:** 10.1101/732800

**Authors:** Cassandra R. Diegel, Steven Hann, Ugur M. Ayturk, Jennifer C.W. Hu, Kyung-eun Lim, Casey J. Droscha, Zachary B. Madaj, Gabrielle E. Foxa, Isaac Izaguirre, VARI Vivarium and Transgenics Core, Noorulain Paracha, Bohdan Pidhaynyy, Terry L. Dowd, Alexander G. Robling, Matthew L. Warman, Bart O. Williams

**Affiliations:** Program in Skeletal Disease and Tumor Microenvironment and Center for Cancer and Cell Biology, Van Andel Research Institute, Grand Rapids, MI 49503; Bioinformatics and Biostatistics Core, Van Andel Research Institute, Grand Rapids, MI 49503; Vivarium and Transgenics Core, Van Andel Research Institute, Grand Rapids, MI 49503; Orthopedic Research Labs, Boston Children’s Hospital and Department of Genetics, Harvard Medical School, Boston, MA 02115; Musculoskeletal Integrity Program, Hospital for Special Surgery Research Institute, New York, NY 10021; Department of Anatomy and Cell Biology, Indiana University School of Medicine, Indianapolis, IN 46202; Department of Chemistry, Brooklyn College, Brooklyn, NY; Department of Biology, Brooklyn College, Brooklyn, NY; Ph.D. Program in Chemistry and Ph.D. Program in Biochemistry, The Graduate Center of the City University of New York, New York, NY 10016

## Abstract

Osteocalcin (OCN), the most abundant non-collagenous protein in the bone matrix, is reported to be a bone-derived endocrine hormone with wide-ranging effects on many aspects of physiology, including glucose metabolism and male fertility. Many of these observations were made using an OCN-deficient mouse allele (Osc^-^) in which the 2 OCN-encoding genes in mice, *Bglap* and *Bglap2*, were deleted in ES cells by homologous recombination. Here we describe mice with a new *Bglap* and *Bglap2* double knockout (dko) allele (Bglap/2^p.Pro25fs17Ter^) that was generated by CRISPR/Cas9-mediated gene editing. Mice homozygous for this new allele do not express full length *Bglap* or *Bglap2* mRNA and have no immunodetectable OCN in their plasma. FTIR imaging of cortical and trabecular bone in these homozygous knockout animals finds alterations in the crystal size and maturity of the bone mineral, hydroxyapatite, compared to wild-type littermates; however, μCT and 3-point bending tests do not find differences from wild-type littermates with respect to bone mass and strength. In contrast to the previously reported OCN-deficient mice with the Osc^-^ allele, blood glucose levels and male fertility in the OCN-deficient mice with *Bglap/2*^pPro25fs17Ter^ allele did not have significant differences from wild-type littermates. We cannot explain the absence of endocrine effects in mice with this new knockout allele. Potential explanations include effects of each mutated allele on the transcription of neighboring genes, and differences in genetic background and environment. So that our findings can be confirmed and extended by other interested investigators, we are donating this new *Bglap* and *Bglap2* double knockout strain to The Jackson Laboratory for academic distribution.

**Author Summary:** Cells that make and maintain bone express proteins that function locally or systemically. The former proteins, such as type 1 collagen, affect the material properties of the skeleton while the latter proteins, such as fibroblast growth factor 23, enable the skeleton to communicate with other organ systems. Mutations that affect the functions of most bone cell expressed proteins cause diseases that have similar features in humans and other mammals, such as mice; for example, brittle bone diseases for type 1 collagen mutations and hypophosphatemic rickets for fibroblast growth factor 23 mutations.

Our study focuses on another bone cell expressed protein, osteocalcin, which has been suggested to function locally to affect bone strength <u>and</u> systemically as hormone. Studies using osteocalcin knockout mice led other investigators to suggest endocrine roles for osteocalcin in regulating blood glucose levels, male fertility, muscle mass, brain development, behavior and cognition. We therefore decided to generate a new strain of osteocalcin knockout mice that could also be used to investigate these non-skeletal effects.

To our surprise the osteocalcin knockout mice we created do not significantly differ from wild-type mice for the 3 phenotypes we examined: bone strength, blood glucose levels, and male fertility. Our data are consistent with findings from osteocalcin knockout rats, but inconsistent with data from the original osteocalcin knockout mice. Because we do not know why our new strain of osteocalcin knockout mice fails to recapitulate phenotypes previously reported for another knockout mouse stain, we have donated our mice to a public repository so that they can be easily obtained and studied in other academic laboratories.

## Introduction

Osteocalcin (OCN) is a protein almost exclusively expressed by osteoblasts [1]. Once transcribed and translated, the 95 amino acid OCN prepromolecule is cleaved to produce a biologically active 46 amino acid, carboxyl-terminal, fragment that contains 3 γ-carboxyglutamic acid residues (residues 62, 66, 69 in the mouse prepromolecule) which are made post-translationally via a vitamin K dependent process [2]. The binding of Ca^2+^ to these γ-carboxyglutamic acid residues causes conformational changes that increase OCN binding to the bone mineral calcium hydroxyapatite [3]. During bone resorption the acidic pH within the bone resorption pit decarboxylates OCN, enabling uncarboxylated OCN release into the bloodstream [4]. Uncarboxylated OCN may also be directly released into the bloodstream by osteoblasts.

In humans OCN is encoded by a single bone gamma-carboxyglutamic acid-containing protein gene (*BGLAP*). In mice OCN in encoded within a 25 kb interval on chromosome 3 that contains *Bglap* and *Bglap2*, (a.k.a., *Og1* and *Og2*), which are highly expressed in bone, and *Bglap3*, (a.k.a., Org), which has minimal expression in bone. The 46 residue carboxyl-terminal domains encoded by *Bglap* and *Bglap2* are 100% identical, and differ by 4 amino acid residues from that encoded by *Bglap3*.

Mice homozygous for a large genomic deletion encompassing *Bglap* and *Bglap2* (*Osc^-^/Osc^-^*) were generated by ES-cell-mediated germline modification and described in 1996 [5]. By 6 months of age, these OCN-deficient mice had qualitatively increased cortical thickness and increased bone density relative to their control littermates. These qualitative differences were associated with significantly increased biomechanical measures of bone strength. Furthermore, OCN-deficient female mice were resistant to oophorectomy-induced bone loss. These data suggested that therapeutic reduction of OCN expression could prevent osteoporosis in humans.

Subsequent studies using *Osc^-^/Osc^-^* mice broadened the biologic role for OCN by identifying wide-ranging physiologic changes when OCN is deficient. The first such report observed that *Osc^-^/Osc^-^* mice had increased visceral fat and displayed elevated blood glucose levels associated with decreased pancreatic beta-cell proliferation and insulin resistance [6]. Subsequent work indicated OCN enhanced male fertility by inducing testosterone production and promoting germ cell survival [7]. Other observed endocrine roles for OCN include influencing fetal brain development and adult animal behavior [8], and promoting adaptation of myofibers to exercise and maintaining muscle mass with ageing [9, 10]. Cumulatively, the reports outlined above with the *Osc^-^/Osc^-^* mice and, subsequently, a conditional knockout strain (Ocn-flox) made by the same investigative team [7] coupled with associated work [11–18] from this team, suggest that therapies which increase uncarboxylated osteocalcin levels may improve glucose intolerance, increase beta islet cell number, reduce insulin resistance, increase testosterone, improve male fertility, enhance muscle mass, and reduce declines in cognition.

Given the potential importance of OCN, we created new strains of OCN-deficient mice using CRISPR/Cas9 gene editing [19]. Specifically, we created mice with mutations that disrupt either *Bglap* or *Bglap2* individually or in combination. Here we report our observations regarding homozygous *Bglap* and *Bglap2* double knockout (Bglap/2^dko/dko^) mice, which we anticipated would recapitulate the previously reported OCN-deficiency phenotypes in the *Osc^-^/Osc^-^* mice of increased bone mass and strength, elevated blood glucose levels, and decreased male fertility. We confirmed that we successfully knocked out *Bglap* and *Bglap2* by performing RNA sequencing and by measuring immunoreactive OCN in mouse plasma. Consistent with previous studies, homozygous offspring with this new OCN-deficient allele (Bglap/2^dko/dko^) were born at the expected Mendelian frequency and exhibited no overt clinical phenotype. Inconsistent with previous studies, we did not observe significantly increased bone mass or strength, significantly elevated blood glucose levels, significantly decreased testosterone levels, or significantly impaired male fertility.

## Results

### Generation of mice with *Bglap and Bglap2* double knockout alleles and OCN-deficiency

By simultaneously injecting Cas9 protein and guide RNAs that recognize sequences within *Bglap* and *Bglap2*, and not within *Bglap3*, we generated several founders that harbor large deletions involving *Bglap* and *Bglap2*. One founder allele, termed p.Pro25fsTer17, is a ~ 6.8 kb deletion that joins exon 2 of *Bglap* to exon 4 of *Bglap2* (Figure 1A). This allele is predicted to produce a chimeric *Bglap/Bglap2* transcript with a reading frame-shift after the 25^th^ amino acid residue and a termination codon 17 residues further downstream. Because p.Pro25 is amino-terminal to the biologically active domain of OCN, mice homozygous for this allele (*Bglap/2*^dko/dko^) should be OCN-deficient.

**Figure 1.**
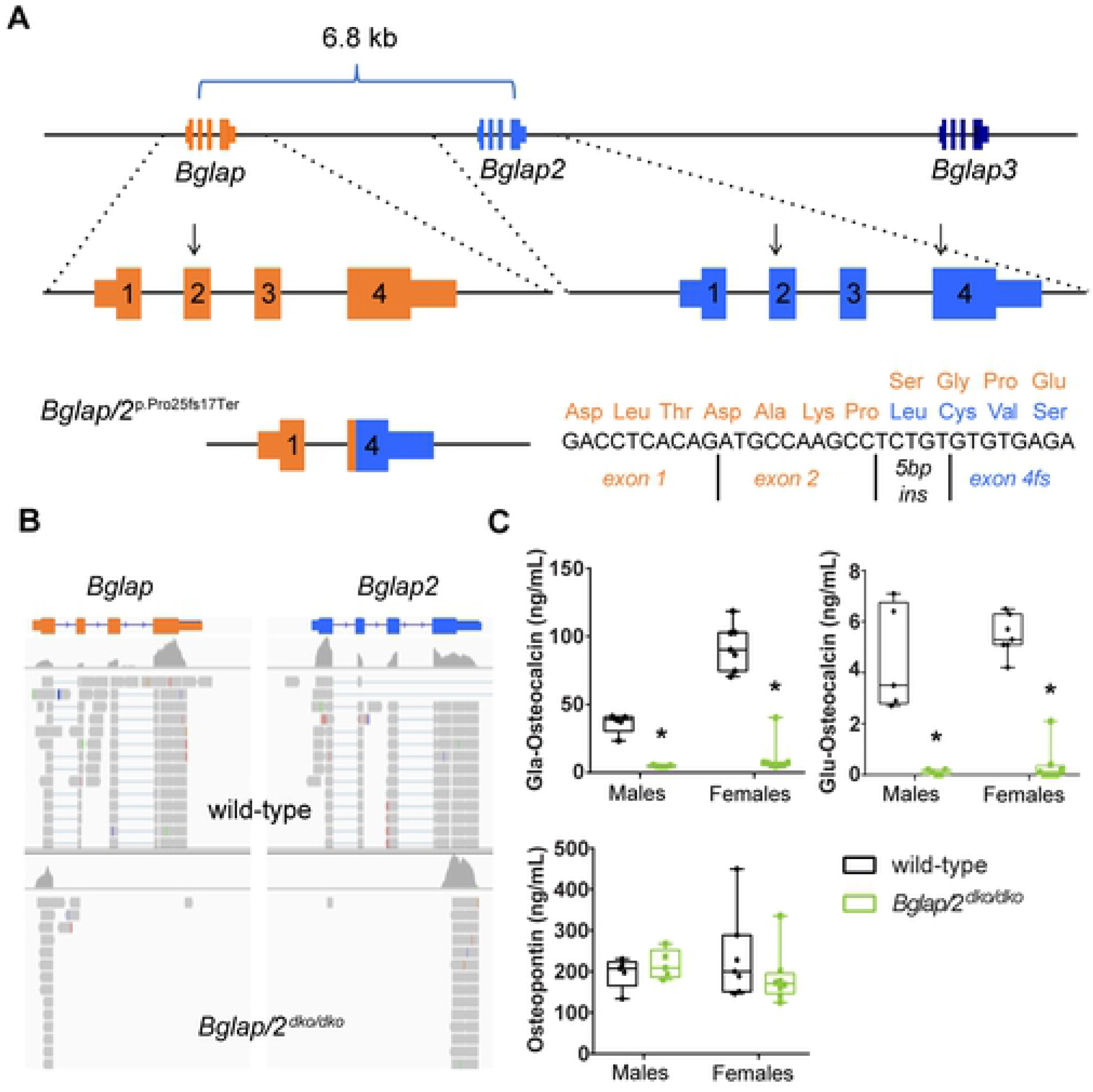
Generation and validation of a *Bglap* and *Bglap2* double knockout allele (*Bglap/2*^dko^) created by CRISPR/Cas9 gene editing. A) Schematic diagram (not drawn to scale) showing the 25 kb interval on mouse chromosome 3 containing *Bglap, Bglap2*, and *Bglap3*. The locations of the guide RNAs used to produce founders with *Bglap* and *Bglap2* intragenic or intergenic knockout alleles are indicated (arrows). The *Bglap/2* double knockout (dko) allele (p.Pro25fs17Ter) deletes a 6.8 kb DNA fragment extending from *Bglap* exon 2 to *Bglap2* exon 4. A 5 bp insertion (5 bp ins) at the DNA ligation site creates a chimeric exon with a reading frame-shift. The frame-shift occurs after p.25 and terminates the protein 17 residues downstream (orange residues indicate wild-type sequence, whereas blue indicates the first 4 frame-shifted residues). B) Integrated Genomics Viewer screenshots of RNA sequencing data from wild-type and homozygous *Bglap* and *Bglap2* double knockout mice (*Bglap/2*^dko/dko^). Note the absence of sequencing reads mapping to *Bglap* exons 2, 3, and 4, and *Bglap2* exons 1, 2, and 3 in the double knockout mice; the mapping algorithm fails to map reads to the *Bglap* chimeric exon 2 because the chimera has a short seed length and a 5 bp insertion. C) Plasma ELISA assays for male and female 6-month-old *Bglap/2*^dko/dko^ (green) and their control (black) littermates for the gamma-carboxyglutamic acid (Gla-OCN) and uncarboxylated (Glu-OCN) forms of OCN as well as Osteopontin (OPN). Each individual mouse is represented as a single dot in each graph. Box and whiskers plots are shown for each measurement with the horizontal line in each representing the median. The following sample sizes were used: male wild-type (n=5), male *Bglap/2*^dko/dko^ (n=5), female wild-type (n=7), and female *Bglap/2*^dko/dko^ (n=8).

*Bglap/2*^dko/dko^ offspring were born from heterozygote crosses at the expected Mendelian frequency and appeared phenotypically indistinguishable from their wild-type and carrier littermates. We confirmed that *Bglap/2*^dko/dko^ homozygous mutants produced the frame-shifted chimeric *Bglap/Bglap2* mRNA transcript, as predicted, by performing RNA sequencing on freshly isolated mouse cortical bone mRNA (Figure 1B). We showed that homozygous mutants were OCN-deficient by measuring plasma carboxylated and uncarboxylated OCN. (Figure 1C). Thus, the p.Pro25fs17 allele eliminates the production of immunodetectable OCN from *Bglap* and *Bglap2*, and, in this respect, is identical to the *Osc^-^* allele [5] and the Cre-excised Ocn-floxed allele [7]. Because a recent report had found that loss of both OCN and Osteopontin was synergistic in terms of effects on bone morphology and mechanical properties [20], we evaluated the plasma levels of Osteopontin. No significant differences in Osteopontin were detected between *Bglap/2*^dko/dko^ mice and their littermate controls (Figure 1C).

### Bglap/2^dko/dko^ mice do not have increased bone mass or strength

Histomorphometric studies detected increased bone area by 6 months of age in *Osc^-^/Osc^-^* mice [5]. Since μCT measures have superseded histomorphometric measures for assessing bone area and bone volume, we collected femurs from 6-month-old *Bglap/2*^dko/dko^ mice and compared their μCT measures to those of their wild-type littermates. We saw no significant differences in cortical or trabecular bone parameters between the *Bglap/2*^dko/dko^ and wild-type mice (Figure 2A-B and Table 1). Femur 4-point bending assays performed on *Osc^-^/Osc^-^* mice had revealed significantly increased bone strength [5]. We performed three-point-bending tests on 6-month-old *Bglap/2*^dko/dko^ and wild-type littermates, and did not find increased bone strength in these OCN-deficient animals (Figure 2C and Supplemental Tables 1 and 2).

**Figure 2.**
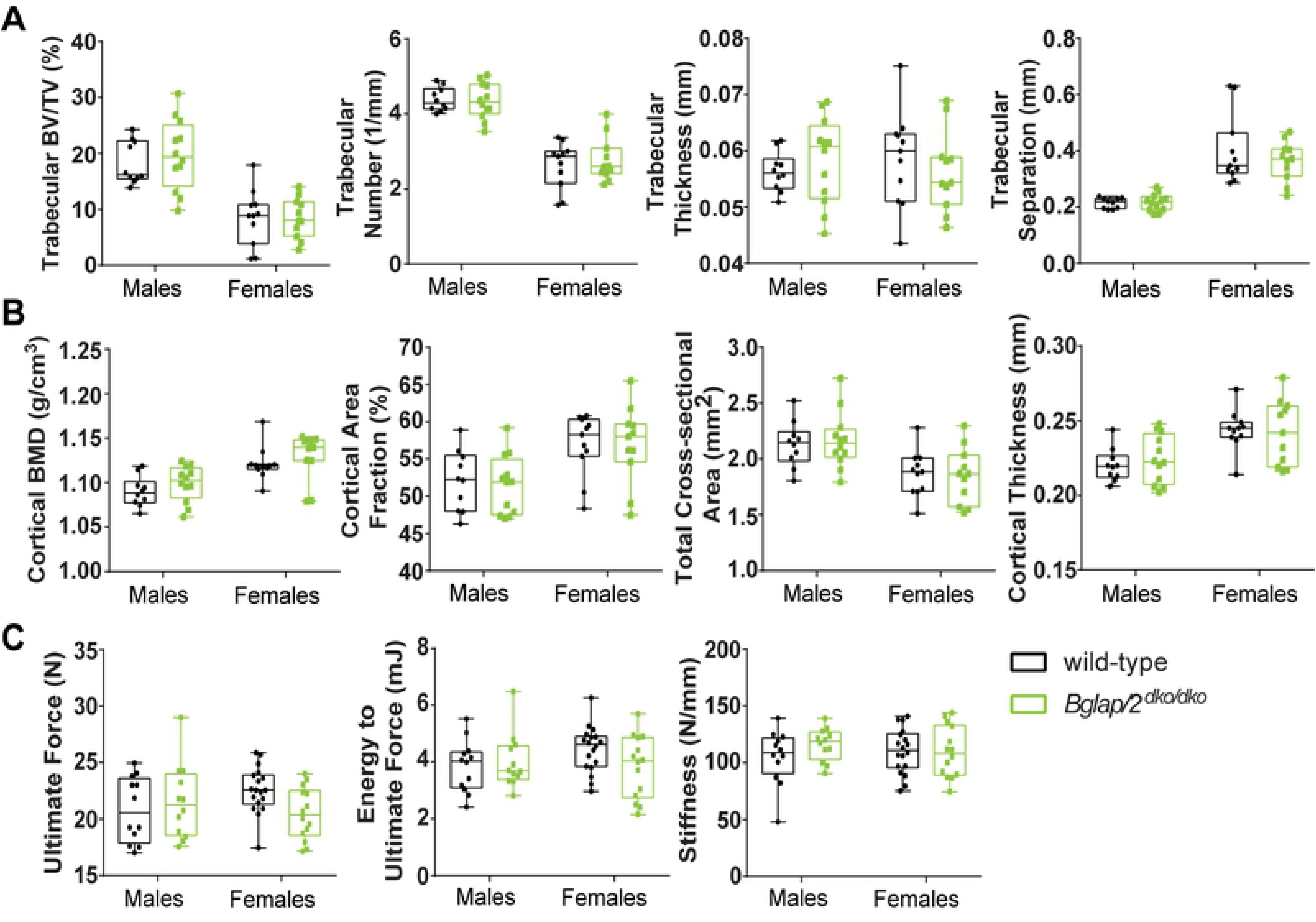
OCN deficiency does not significantly alter bone mass or strength as assessed by μCT and biomechanical testing. A) μCT analysis of trabecular bone parameters detects no significant differences between male or female 6-month-old *Bglap/2*^dko/dko^ (green) and their wild-type (black) littermates. Trabecular bone volume/total volume (BV/TV), trabecular number, thickness, and separation are shown. Additional measurements are included in Table 1. The following sample sizes were used for both cortical and trabecular measurements: male wild-type (n=10), male *Bglap/2*^dko/dko^ (n=12), female wild-type (n=11), and female *Bglap/2*^dko/dko^ (n=11). Box and whiskers plots are shown for each measurement with the horizontal line in each representing the median. μCT analysis of cortical bone parameters detects no significant differences between male or female 6-month-old *Bglap/2*^dko/dko^ (green) and their wild-type (black) littermates. Cortical bone mineral density (BMD), cortical area fraction, tissue area, and cortical thickness are shown as representative measurements. Additional measurements and details are included in Table 1 and sample sizes are noted above. Box and whiskers plots are shown for each measurement with the horizontal line in each representing the median. B) Biomechanical loading assessments detect no significant differences between male or female 6-month-old *Bglap/Bglap2*^dko/dko^ (green) and their wild-type (black) littermates. Ultimate force, energy to ultimate force, and stiffness are shown as representative measurements. Additional measurements and details are included in Supplemental Tables 1 and 2. The following sample sizes were used: male wild-type (12), male *Bglap/2*^dko/dko^ (n=12), female wild-type (n=18), and female *Bglap/2*^dko/dko^ (n=14). Box and whiskers plots are shown for each measurement with the horizontal line in each representing the median.

**Table 1.** uCT analysis of trabecular and cortical bone parameters for male or female 6-month-old *Bglap/2* and their wild-type (*Bglap2*^WT^) littermates. Sample sizes are shown at the top of each column.

| Index | Female |  | Male |  |
| --- | --- | --- | --- | --- |
|  | **Bglap/2<sup>WT</sup> (n=11)** | **Bglap/2<sup>dko/dko</sup> (n=12)** | **Bglap/2<sup>WT</sup> (n=10)** | **Bglap/2<sup>dko/dko</sup> (n=12)** |
| Trab.TV (mm<sup>3</sup>) | 3.181 ± 0.505 | 2.943 ± 0.469 | 3.783 ± 0.375 | 3.835 ± 0.602 |
| Trab.BV (mm<sup>3</sup>) | 0.276 ± 0.174 | 0.258 ± 0.127 | 0.693 ± 0.152 | 0.760 ± 0.249 |
| Trab.BV/TV (%) | 0.085 ± 0.048 | 0.085 ± 0.034 | 0.183 ± 0.036 | 0.198 ± 0.061 |
| Trab.Conn.D. (1/mm<sup>3</sup>) | 32.603 ± 18.917 | 33.891 ± 20.175 | 118.994 ± 22.304 | 112.240 ± 29.851 |
| Trab.SMI | 2.617 ± 0.626 | 2.542 ± 0.441 | 1.779 ± 0.406 | 1.603 ± 0.587 |
| Trab.N (1/mm) | 2.632 ± 0.588 | 2.773 ± 0.557 | 4.386 ± 0.298 | 4.350 ± 0.457 |
| Trab.Th (mm) | 0.059 ± 0.008 | 0.056 ± 0.007 | 0.056 ± 0.003 | 0.058 ± 0.007 |
| Trab.Sp (mm) | 0.401 ± 0.117 | 0.364 ± 0.067 | 0.214 ± 0.018 | 0.217 ± 0.029 |
| Trab.BMC (µgHA/cm<sup>3</sup>) | 0.261 ± 0.170 | 0.239 ± 0.116 | 0.640 ± 0.144 | 0.707 ± 0.240 |
| Cort.TV (mm<sup>2</sup>) | 6.643 ± 0.723 | 6.298 ± 0.635 | 7.059 ± 0.534 | 7.119 ± 0.741 |
| Cort.BV (mm<sup>2</sup>) | 2.910 ± 0.205 | 2.815 ± 0.188 | 2.705 ± 0.170 | 2.740 ± 0.178 |
| Cort.BV/TV (%) | 0.441 ± 0.029 | 0.450 ± 0.036 | 0.384 ± 0.020 | 0.387 ± 0.032 |
| Cort.Th (mm) | 0.244 ± 0.013 | 0.242 ± 0.021 | 0.220 ± 0.011 | 0.224 ± 0.016 |
| Cort.BMC (µgHA/cm<sup>3</sup>) | 3.264 ± 0.256 | 3.181 ± 0.221 | 2.949 ± 0.197 | 3.011 ± 0.204 |
| Cort.BMD (µgHA/cm<sup>3</sup>) | 1.121 ± 0.018 | 1.130 ± 0.025 | 1.090 ± 0.016 | 1.099 ± 0.019 |
| CAF (%) | 0.569 ± 0.039 | 0.570 ± 0.050 | 0.521 ± 0.039 | 0.515 ± 0.038 |
| T.Ar (mm) | 1.870 ± 0.193 | 1.848 ± 0.239 | 2.123 ± 0.195 | 2.166 ± 0.241 |
| Cs.Th (mm) | 0.244 ± 0.013 | 0.242 ± 0.021 | 0.220 ± 0.011 | 0.224 ± 0.016 |
| Data represents mean ± SD |  |  |  |  |

### Fourier-transform infrared imaging (FTIR) reveals increased bone crystal size and mineral maturity in *Bglap/2^dko/dko^* mice

To gain additional insight into the constitution of the bone matrix in *Bglap/2*^dko/dko^ animals, we evaluated data from FTIR images collected in cortical and trabecular bone sections (Figure 3 and Supplemental Table 3). There was a significant increase (p < 0.001) in mineral crystallinity of cortical bone in the *Bglap/2*^dko/dko^ mice compared to their control littermates. The crystallinity parameter is related to bone crystal size and how close the mineral is to hydroxyapatite stoichiometry (Ca_10_ (PO_4_)_6_ (OH)_2_). Both carbonate and acid phosphate (HPO4^2-^) ion can incorporate into the hydroxyapatite lattice and acid phosphate is observed in new mineral deposition [21]. There was a significant (p < 0.01) decrease in the carbonate to mineral ratio and in acid phosphate substitution (p < 0.02) in the *Bglap/2*^dko/dko^ bones. Trabecular bone showed a similar increase in crystallinity and decrease in acid phosphate indicating increased crystal size and greater mineral maturity in the OCN-deficient bones.

**Figure 3.**
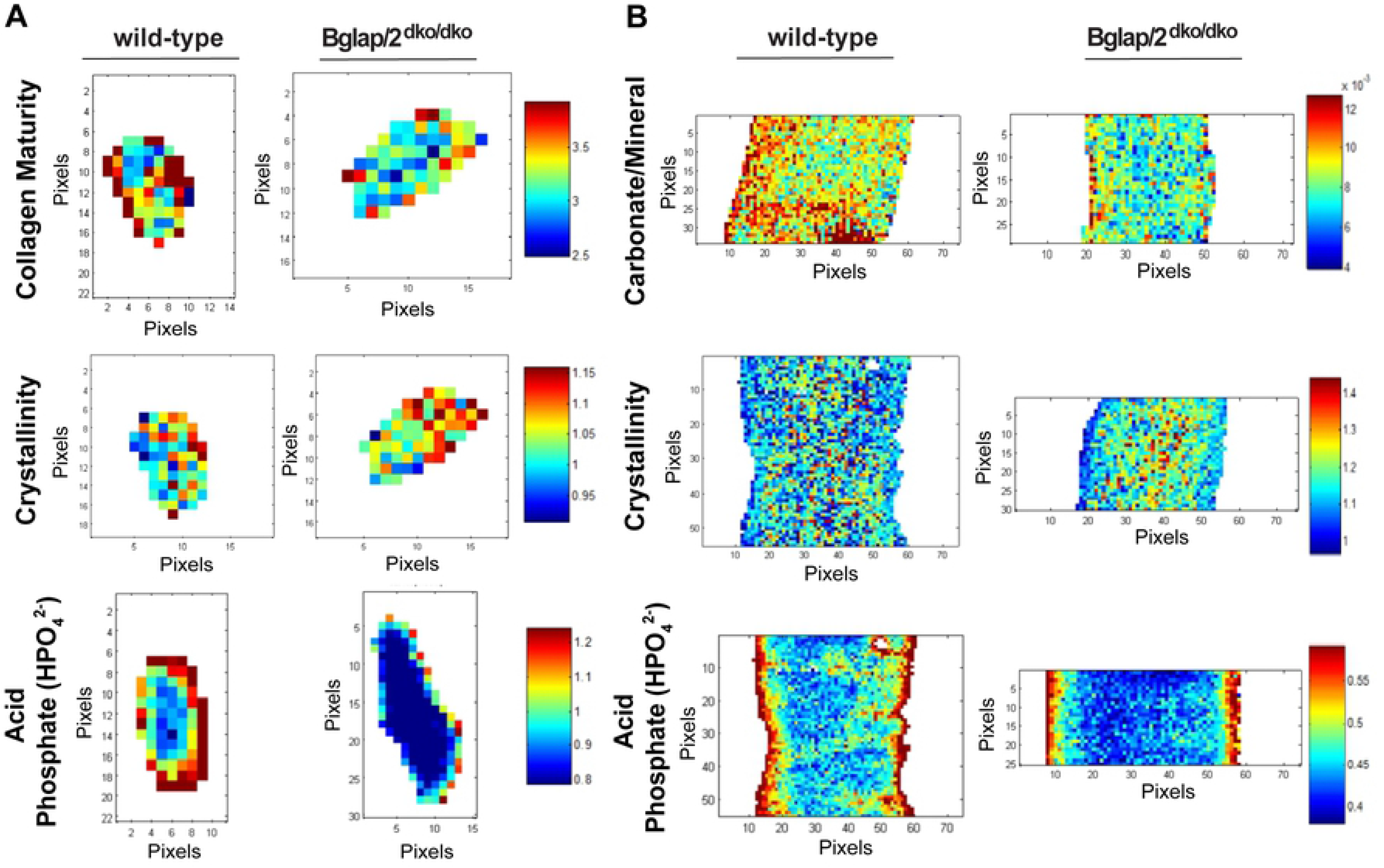
Fourier-transform infrared imaging (FTIR) reveals differences in calcium hydroxyapatite crystal size and maturity between *Bglap/2^dko/dko^* and wild-type mice. A) FTIR images of trabecular bone showing the spatial distribution of the parameters for wild-type and *Bglap/2*^dko/dko^ female mice. Representative images show collagen crosslink maturity is decreased in the *Bglap/2*^dko/dko^ mineral, crystallinity is increased in the *Bglap/2*^dko/dko^ mineral and acid phosphate (HPO_4_^2-^) is decreased in the *Bglap/2*^dko/dko^ mineral. Additional measurements and details are included in Supplemental Table 3. B) FTIR images of cortical bone showing the spatial distribution of the parameters wild-type and *Bglap/2*^dko/dko^ female mice. Representative images show carbonate//mineral ratio is decreased in the *Bglap/2*^dko/dko^ mineral, the crystallinity is increased in the *Bglap/2*^dko/dko^ mineral and the acid phosphate (HPO_4_^2-^) content is decreased in the *Bglap/2*^dko/dko^ mineral. Additional measurements and details are included in Supplemental Table 3.

### *Bglap/2^dko/dko^* mice have normal blood glucose levels

The first reported endocrinologic role for OCN involved regulating glucose levels. From 1 to 6 months of age *Osc^-^/Osc^-^* mice consistently exhibited increased random blood glucose levels compared to controls (p < 0.05) [6]. Fasting blood glucose levels were also increased in 6-month-old *Osc^-^/Osc^-^* mice compared to controls (p < 0.001) [4, 6]. We observed no differences in blood glucose levels between 5 to 6-month-old *Bglap/2*^dko/dko^ mice and their wild-type littermates, when measured either after an overnight fast or at midday in animals with *ad libitum* access to food (Figure 4). We also observed no difference in weight between 6-month-old *Bglap/2*^dko/dko^ and wild-type mice (Figure 4).

**Figure 4.**
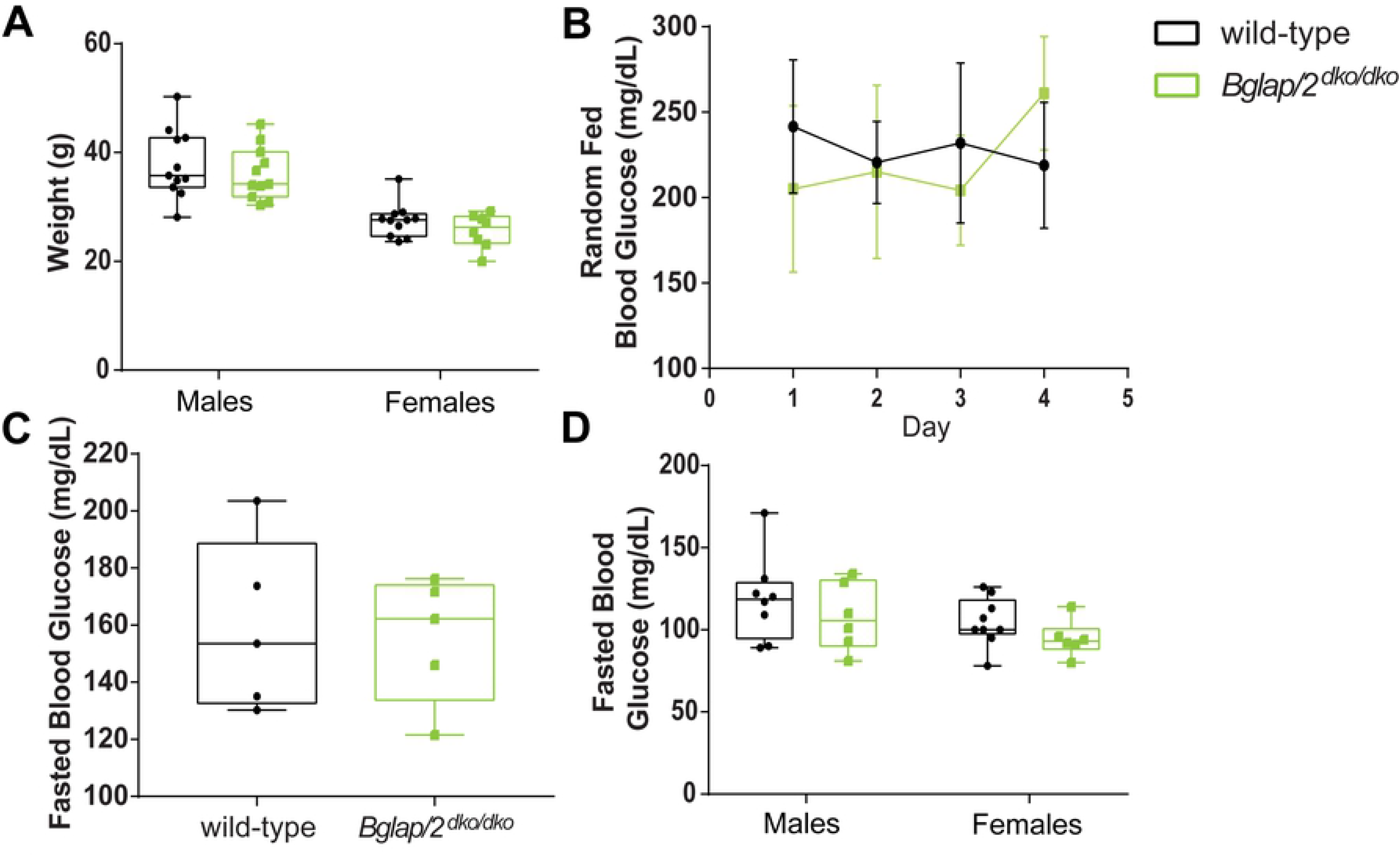
No significant evidence (all p values >0.05) that whole body weight, random fed glucose levels or fasting glucose values differ between *Bglap/2*^dko/dko^ and wild-type mice. A) Total weight for 6-month old *Bglap/2*^dko/dko^ and wild type males and females is shown. Sample sizes are male wild-type (n=11), male *Bglap/2*^dko/dko^ (n=11), female wild-type (n=11), and female *Bglap/2*^dko/dko^ (n=8). Box and whiskers plots are shown for each measurement with the horizontal line in each representing the median. B) Five wild-type and 5 *Bglap/2*^dko/dko^ female 5 to 6 month-old mice were sampled for blood glucose levels on 4 consecutive days 6 hours into their light cycle while having *ad libitum* access to food. At least 2 glucose measurements were taken on each day for each mouse. The average of these daily readings for each mouse was calculated and then the means for each of the genotypic cohorts was calculated and is presented. Means and standard deviations are shown. C) The animals in 4B were also fasted for 16 hours before glucose levels were assessed. This was done on 2 occasions approximately 1 week apart. At least 2 glucose measurements were taken on each day for each mouse. The average of these daily readings for each mouse was calculated and is presented as box and whiskers plots with the horizontal line in each representing the median. In contrast to the data presented in 4D, these mice were not euthanized at the time of glucose assessment. D) Blood glucose levels were measured in 6-month-old *Bglap/2*^dko/dko^ and wild-type males and females after an overnight fast at the time of euthanasia. Sample sizes are male wild-type (n=11), male *Bglap/2*^dko/dko^ (n=11), female wild-type (n=11), and female *Bglap/2*^dko/dko^ (n=8). Box and whiskers plots are shown for each measurement with the horizontal line in each representing the median.

### *Bglap/2^dko/dko^* male mice have normal fertility

The second reported endocrinologic role for OCN involved male fertility. *Osc^-^/Osc^-^* male mice had smaller testes, lower testosterone levels, and produced fewer and smaller litters than control mice [7, 17]. We therefore measured testis size, testosterone levels, and litter size in male *Bglap/2*^dko/dko^ and wild-type littermate mice. We found no significant differences in any of these measures between *Bglap/2*^dko/dko^ and wild-type mice (Figure 5).

**Figure 5.**
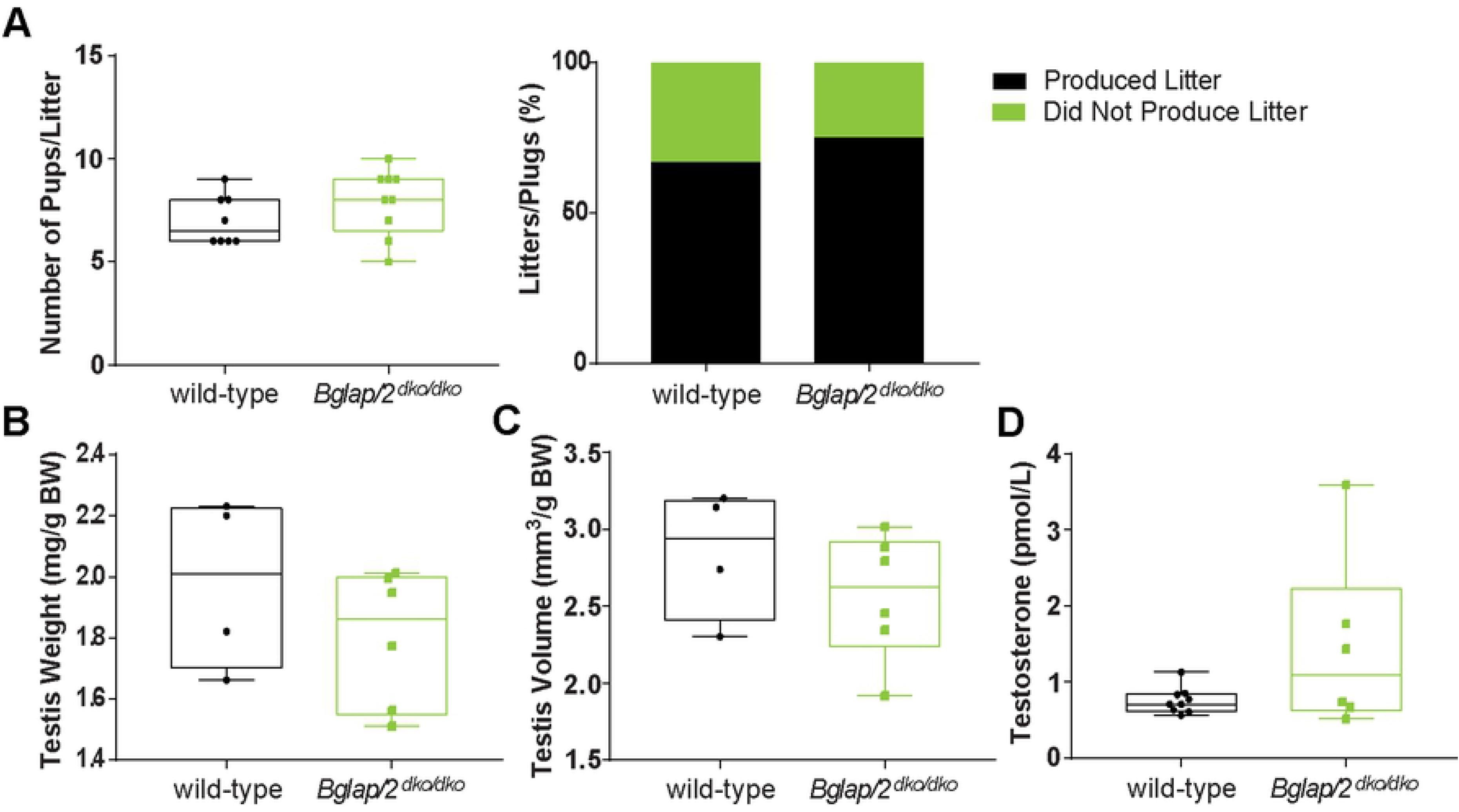
No significant evidence of fertility being affected *Bglap/2*^dko/dko^ mice (all p values >0.05). A) The number of pups per litter and the percent conception rate (n=11 for wild-type and n=12 for *Bglap/2*^dko/dko^ total matings for each genotype) resulting from crosses of *Bglap/2*^dko/dko^ males or their wild-type littermates to wild-type C57Bl/6J females are shown. Box and whiskers plots are shown for number of pups/litter with the horizontal line in each representing the median. B) Testis weight as expressed as mg dry testis weight/gram of total body weight for wild-type (n=4) and *Bglap/2*^dko/dko^ (n=6) males is shown. Box and whiskers plots are shown for each measurement with the horizontal line in each representing the median. C) Testis volume as expressed as mm^3^/gram of total body weight for wild-type (n=4) and *Bglap/2*^dko/dko^ (n=6) males is shown. Box and whiskers plots are shown for each measurement with the horizontal line in each representing the median. D) Plasma testosterone levels from 6-month-old *Bglap/2*^dko/dko^ males (n=6) and wild-type (n=9) littermates are shown.

### Few differences in cortical bone mRNA expression between *Bglap/2^dko/dko^* and control mice

RNA sequencing of cortical bone samples from *Bglap/2^dko/dko^* mice and their wild-type littermates confirmed the absence of wild-type *Bglap* and *Bglap2* transcripts (Figure 1B). Although we sequenced a limited number of specimens from each group, we also performed a transcriptome-wide differential expression analysis searching for large transcriptional changes in bone tissue. We generated RNA sequencing libraries from 4-month-old male mice that on average yielded 46 million reads and 82% uniquely mapping reads. After performing *in silico* filtering to remove contaminating blood, marrow, and muscle transcripts, we identified 14 transcripts that differed significantly between the OCN-deficient and wild-type mice after correcting for multiple hypothesis testing, and which had an average FPKM >3 in mice with either or both genotypes (Table 2 and Supplemental Table 4). As expected, the mean FPKM for *Bglap* was reduced from 2157 in wild-type mice to 158 in *Bglap/2*^dko/dko^ mice. The mean FPKM for *Bglap2* did not significantly differ between wild-type and *Bglap/2*^dko/dko^ mice (1070 versus 1310 and Supplemental Table 4), with most reads in the mutant mice mapping to exon 4. Thus, despite the chimeric *Bglap/2* mRNA containing a frame-shift and premature truncation codon it is not subject to nonsense-mediated mRNA decay. The low level of *Bglap3* transcript (mean FPKM 10) seen in wild-type cortical bone increased 8-fold (mean FPKM 79, adjusted p < 0.001) in *Bglap/2*^dko/dko^ bone remains less than 2.5% of the combined FPKMs for *Bglap* and *Bglap2* in wild-type bone. It is also notable that 6 of the remaining 11 transcripts in Table 2 are encoded on chromosome 3, raising the possibility that CRISPR/Cas9 gene editing altered regulatory elements that directly affect these genes. Similar off target mechanisms could explain the altered expression of transcripts encoded by other chromosomes. Alternatively, these gene expression changes could be the result of OCN-deficiency. Although our RNA sequencing data might be underpowered to detect small changes (i.e., 2-fold or less) in transcript abundance, we found no significant differences in the expression of osteoblast and osteocyte transcripts, including *Alpl, Col1a1, Col1a2, Runx2, Opn, Sost, Dmp1*, and *Fgf23*.

**Table 2.** mRNA Transcripts with statistically significant differences in transcript abundance between *Bglap*/2^dko/dko^ and wild-type mice. Mean FPKMs for each cohort are shown, as are the genomic coordinates for each gene. Note that in addition to *Bglap* and *Bglap3*, 6 of the remaining 11 genes also map to chromosome 3.

| Ensembl ID | Gene ID | p-value<br>adj | Mean<br>dko/dko<br>FPKM | Mean<br>WT<br>FPKM | Chr Start and End |
| --- | --- | --- | --- | --- | --- |
| ENSMUSG00000067017 | *Gm3608* | 3.22E-30 | 11.7 | 0.7 | 1; 105455971 - 105458829 |
| ENSMUSG00000074483 | *Bglap* | 5.38E-23 | 158.3 | 2156.7 | 3; 88383501 - 88384464 |
| ENSMUSG00000028081 | *Rps3a1* | 2.56E-11 | 65.2 | 287.1 | 3; 86137940 - 86142702 |
| ENSMUSG00000074340 | *Ovgp1* | 2.42E-06 | 4.1 | 1.0 | 3; 105973711 - 105987423 |
| ENSMUSG00000091383 | *Hist1h2al* | 5.55E-05 | 23.9 | 0.1 | 13; 51202727 - 51203065 |
| ENSMUSG00000040809 | *Chil3* | 0.000414 | 51.5 | 247.7 | 3; 106147554 - 106167564 |
| ENSMUSG00000074489 | *Bglap3* | 0.000477 | 77.5 | 9.8 | 3; 88368616 - 88372743 |
| ENSMUSG00000106508 | *4933425M03Rik* | 0.004872 | 4.0 | 0.9 | 3; 85887691 - 85889234 |
| ENSMUSG00000076564 | *Igkv12-46* | 0.005734 | 5.9 | 25.1 | 6; 69764523 - 69765022 |
| ENSMUSG00000068855 | *Hist2h2ac* | 0.009698 | 0.3 | 3.5 | 3; 96220361 - 96220880 |
| ENSMUSG00000051906 | *Cd209f* | 0.010412 | 3.2 | 0.8 | 8; 4102787 - 4105728 |
| ENSMUSG00000028104 | *Polr3gl* | 0.015468 | 50.0 | 22.5 | 3; 96577872 - 96594181 |
| ENSMUSG00000017861 | *Mybl2* | 0.019515 | 2.0 | 4.2 | 2; 163054687 - 163084688 |

## Discussion

We created a double knockout allele for *Bglap* and *Bglap2* (*Bglap/2*^dko^) and then evaluated bone properties, glucose levels, and male fertility in homozygous mutant mice and their wild-type littermate controls. We only observed skeletal differences between *Bglap/2*^dko/dko^ and wild-type littermates using FTIR. These differences revealed an increase in crystallinity in the *Bglap/2*^dko/dko^ mice, consistent with the previously reported role for OCN in regulating bone crystal size [22]. We also found more mature mineral in the *Bglap/2*^dko/dko^ mice suggesting osteocalcin could be involved in mineral maturation.

In contrast to prior reports that utilized global double knockout [5] or conditional double knockout alleles [7], we did not observe any significant effect of OCN deficiency on bone mass and strength (Figure 2), blood glucose levels and body weight (Figure 4), and male fertility (Figure 5). Lee et al [6] observed that OCN-deficient *Osc^-^/Osc^-^* mice had significantly increased random fed blood glucose levels at 1, 3, and 6 months of age, and significantly increased overnight fasted blood glucose levels at 6 months of age. We found no difference in blood glucose levels between *Bglap/2*^dko/dko^ mice and their wild-type littermates. Oury *et al*. [7] observed male *Osc^-^/Osc^-^* mice had significantly lower litter frequencies and sizes, testicular weights and volumes, and plasma testosterone levels. We did not observe significant effects in the *Bglap/2*^dko/dko^ mice.

We do not know why *Bglap/2*^dko/dko^ mice do not replicate the endocrinologic phenotypes that were attributed to other *Bglap* and *Bglap2* double knockout mice [4, 6, 7, 17]. Our RNA sequencing indicates we disrupted both *Bglap* and *Bglap2* transcripts (Figure 1B) and our plasma ELISA data indicate that OCN is undetectable in *Bglap/2*^dko/dko^ mice. We cannot preclude the possibility that increased *Bglap3* mRNA expression compensates for the absence of *Bglap* and *Bglap2* in the *Bglap/2*^dko/dko^ mice. Although we could not detect OCN by ELISA in *Bglap/2*^dko/dko^ animals, the *Bglap3* protein product may fail to be detected with this assay since it differs by 4 amino acid residues from that produced by *Bglap* and *Bglap2*. However, we think compensation by *Bglap3* is unlikely since its cortical bone transcript abundance in the *Bglap/2*^dko/dko^ is less than 2.5% of the *Bglap* and *Bglap2* transcript abundance in wild-type mice.

At present we do not know whether the increase in *Bglap3* mRNA expression is compensatory or an artifact of moving the *Bglap* promoter 6.8 kb nearer to *Bglap3*. The *Bglap* promoter appears to have been deleted along with *Bglap* and *Bglap2* in the *Osc^-^/Osc^-^* mice [5], but not in the *Ocn-flox* conditional knockout mice [7]. Therefore, comparing RNA sequencing data across all 3 strains could identify allele-specific effects on gene expression that account for phenotype differences between the *Bglap/2*^dko/dko^ mice and the previously published mice. Other explanations for differences in phenotype could include genetic background, vivarium environment, genetic and epigenetic changes across generations, and assay design.

Although we did not observe phenotypes previously ascribed to OCN-deficiency in mice, our data do align with those reported for the OCN knockout rat. Rats, like humans, have only a single *Bglap* locus. *Bglap* knockout rats do not have elevated glucose levels, insulin resistance, or decreased male fertility [23]. To date, genetic studies in humans have also not identified a role for OCN in these aspects of physiology. No human Mendelian genetic disease has yet been attributed to loss-of-function mutations in BGLAP, in the Online Mendelian Inheritance in Man or MatchMaker exchange databases [[24, 25] https://omim.org and https://www.matchmakerexchange.org, each accessed on August 5, 2019], and there seems to be tolerance for heterozygous loss-of-function mutations at the population level (pLI = 0) [26] in the gnomAD database [https://gnomad.broadinstitute.org]. Two putative loss of function mutations (p.Gly27Ter19/rs1251034119 and p.Tyr52Ter/rs201282254) have carrier frequencies in the genome aggregation datable (gnomAD) of 1-in-120 in African and ~ 1-in-150 in Ashkenazi Jewish participants, respectively. Also, 1 African participant and 1 Ashkenazi Jewish participant is homozygous for loss-of-function mutations in gnomAD. There is also no evidence from Genome-Wide Association Studies that common variants near *BGLAP* influence bone mineral density, blood glucose levels, body mass index, or risks for developing diabetes, autism, or psychiatric disease [https://www.gwascentral.org, http://pheweb.sph.umich.edu, http://www.type2diabetesgenetics.org]. Mutations in *BGLAP* also do not appear to be enriched among children with autism or severe intellectual disability who have undergone research sequencing [BioRvix: https://doi.org/10.1101/484113].

Because we did not find abnormalities in the *Bglap/2*^dko/dko^ mice that have been reported for the *Osc^-^/Osc^-^* mice, we chose not to perform studies interrogating the role of OCN on muscle mass, central nervous system development, and behavior. Instead, we are donating our mice to The Jackson Laboratory (JAX stock # 032497, allele symbol Del(3Bglap2-Bglap)1Vari). We suggest the *Osc^-^/Osc^-^* and Ocn-flox mice also be donated to a resource that facilitates public distribution, so interested investigators can identify why different *Bglap* and *Bglap2* double knockout alleles produce different phenotypes.

## Materials and Methods

### Experimental Animals

Mice were maintained in accordance with institutional animal care and use guidelines, and experimental protocols were approved by the Institutional Animal Care and Use Committee of the Van Andel Institute. These animals are available from The Jackson Laboratory (Stock number 032497).

### Generation of *Bglap/Bglap2* deletion mice using CRISPR/Cas9

Alterations in the mouse *Bglap and Bglap2* alleles were created using a modified CRISPR/Cas9 protocol [27]. Briefly, two sgRNAs targeting exon 2 of OG1(AGACTCAGGGCCGCTGGGCT) and exon 4 of OG2 (GGGATCTGGGCTGGGGACTG) were designed using MIT’s guide sequence generator (crispr.mit.edu.). The guide sequence was then cloned into vector pX330-U6-Chimeric_BB-CBh-hSpCas9 which was a gift from Feng Zhang (Addgene plasmid # 42230; http://n2t.net/addgene:42230; RRID:Addgene_42230). The T7 promoter was added to the sgRNA template, and the sequence was synthetized by IDT. The PCR amplified T7-sgRNA product was used as template for *in vitro* transcription using the MEGAshortscript T7 kit (Thermo Fisher Scientific). The injection mix consisted of Cas9 mRNA (Sigma Aldrich) (final concentration of 50 ng/ul) and sgRNA’s (20 ng/ul) in injection buffer (10 mM Tris; 0.1 mM EDTA pH 7.5) injected into the pronucleus of C57BL/6;C3H zygotes. After identifying founders we backcrossed the line to C57BL/6 twice before intercrossing to generate animals for our study.

### Genotypic Identification

To genotype the *Bglap/2*^dko^ allele we used the following primers: OG1-E1-Fwd (ACACCATGAGGACCATCTTTC) and OG1-E4-Rev (AGGTCATAGAGACCACTCCAGC) to amplify a 517bp wildtype product and in a separate reaction we used OG1-E1-Fwd and OG1(E2)-OG2(E4)-Rev (AAGCTCACACACAGAGGCTTGG) to amplify a 238bp *Bglap/2*^dko^ allele product.

### Blood Chemistry Analysis

Blood was harvested into an EDTA-treated tube at the time of euthanasia via intracardiac puncture, spun down at 8,000 rpm for 6 minutes, and plasma collected and stored at −80°C until use. Enzyme immunoassays were used to measure plasma concentrations of mouse Glu-Osteocalcin (MK129;Takara), mouse Gla-Osteocalcin (MK127; Takara), mouse Osteopontin (MOST00; R&D Systems), and mouse Testosterone (55-TESMS-E01; Alpco) according to the manufacturers’ recommendations. Statistical analysis was performed used a linear regression model to test for differences between genotypes via R.

### Collection of Samples for Skeletal Analysis

Femurs were collected at 26 weeks of age; the right hindlimb was fixed in neutral buffered formalin for μCT analysis and the left hindlimb was frozen in saline-saturated gauze for biomechanical testing.

### μCT and Mechanical Testing

Formalin-fixed femora were scanned, reconstructed, and analyzed as previously described [28]. Briefly, 10 μm resolution, 50-kV peak tube potential, 151-ms integration time, and 180° projection area were used to collect scans on a Scanco μCT-35 desktop tomographer. The distal 60% of each femur was scanned, thresholded, and reconstructed to the 3^rd^ dimension using Scanco software. Standard parameters related to cancellous and cortical bone mass, geometry, and architecture were measured [29].

For evaluation of bone mechanical properties, frozen femur samples were brought to room temperature over a 4 hr period, then mounted across the lower supports (8 mm span) of a 3-point bending platen, mounted in a TestResources R100 small force testing machine. The samples were tested in monotonic bending to failure using a crosshead speed of 0.05 mm/sec. Parameters related to whole bone strength were measured from force/displacement curves as previously described [30, 31].

### Fourier-transform infrared imaging (FTIR) Analysis

Femora from female, 6-month old wild-type and *Bglap/2^dko/dko^* mice were cleaned of soft tissue, processed and embedded in polymethylmethacrylate (PMMA) [32]. Longitudinal sections, 2 μm thick, were mounted on infrared windows where spectral images were collected at a 4 cm^-1^ spectral resolution and ~7 μm spatial resolution from a Spotlight 400 Imaging system (Perkin Elmer Instruments, Shelton, CT USA). Background spectra were collected under identical conditions from clear Ba_2_F windows and subtracted from sample data by instrumental software. IR spectra were collected from three areas (~ 500 μm x 500 μm) of cortical bone per sample. The spectra were baseline-corrected, normalized to the PMMA peak at 1728 cm^-1^and the spectral contribution of PMMA embedding media was subtracted using ISYS Chemical Imaging Software (Malvern, Worcestershire, UK). Spectroscopic parameters of carbonate-to-mineral ratio, crystallinity, mineral-to-matrix ratio, collagen maturity and acid phosphate were calculated. The carbonate to mineral ratio is the integrated area ratio of the carbonate peak (850-890)/ν_1_ ν_3_ PO_4_ band (900-1200 cm^-1^) while the mineral-to-(collagen)-matrix ratio is the integrated area ratio of the ν_1_ ν_3_ PO_4_ band (900-1200 cm^-1^) / amide I band (1590-1712 cm^-1^). The mineral crystallinity parameter corresponds to the crystallite size and perfection as determined by x-ray diffraction and is calculated from the intensity ratios of subbands at 1030 cm^-1^ (stoichiometric apatite) and 1020 cm^-1^ (nonstoichiometric apatite). The collagen maturity parameter is the ratio of nonreducible (mature) to reducible (immature) collagen cross-links, which is expressed as the intensity ratio of 1660 cm^-1^ /1690 cm^-1^. The acid phosphate content in the mineral is measured from the peak height ratio of 1128/1096. The result for each parameter was reported as a histogram, describing the pixel distribution in the image. The mean value of the distribution was reported and associated color-coded images were generated at the same time by ISYS. Data for each measured parameter are expressed as mean ± standard error of the mean for each group. The data for all measured parameters were found to be normally distributed as analyzed by the Shapiro-Wilk tests did not find enough evidence to conclude any measured parameters were non-normally distributed. The average values were compared by the Student’s independent t-test for significant differences between groups. Differences for each measured parameter were considered statistically significant when p ≤ 0.05.

### Baseline and random glucose measurements

Cohorts of 5-6 month-old female mice, wild-type (n=5) and *Bglap/2^dko/dko^* (n=5), were subjected to 4 random glucose measurements taken 6 hours into their light cycle by tail nick using a glucose meter (AlphaTRAK; Zoetis) on 4 consecutive days. These same cohorts had glucose measurements obtained by tail nick after an overnight fast during which the animals had access to water on two occasions one week apart.

Additional cohorts of mice were fasted overnight, but with access to water, prior to euthanasia by CO2 inhalation. Glucose levels were measured immediately after euthanasia. Animals were euthanized by CO_2_ inhalation and glucose levels were immediately measured from blood collected by tail nick using a glucose meter.

Glucose data were analyzed using a linear mixed-effects model via the R package *lme4* [33] to account for repeated sampling. Normality of the residuals was verified visually using a qq-plot.

### Measurement of testis size and weight

Testes were removed from 6-month old males after euthanasia and fixation. Fatty tissue was carefully removed from each testis and the sample dried prior to weighing on an analytical scale to determine dry weight. Testis weight was normalized to body weight (mg/g BW). Each testis was imaged and the length and width calculated using Nikon’s NIS-Elements documentation software. Testis volume was normalized to body weight (mm^3^/g BW). Statistical analysis was performed using a linear regression model to test for differences between genotypes via R.

### Assessment of Fertility

Male mice of mating age (7-16 weeks of age) were singly housed for 1 week before mating. Males were mated to C57BL/6J females in the evening and vaginal plugs checked the following morning. Number of pups per litter were compared between genotypes and the percentage of vaginal plugs resulting in delivery and the number of pups per delivery were noted. Logistic mixed-effects regression with a random intercept for paternal mouse was used to determine if the rate of conception differed significantly between the two genotypes. A poisson mixed-effects model with a random intercept for paternal mouse was used to determine if the number of pups per delivery differed significantly between the two genotypes. Both models were analyzed via the R v 3.5.2 (https://cran.r-project.org/) package *lme4* [33]. Methods for testosterone measurement are provided in the previous section on blood chemistry analysis.

### Cortical bone RNA sequencing and data analysis

Tibial cortical bone was recovered from 4-month-old male wild-type (n=3) and *Bglap/2*^dko/dko^ (n=5) mice immediately following euthanasia by CO2 inhalation as previously described [34]. Samples were frozen in liquid nitrogen, pulverized, and suspended in TRIZol. Total RNA was recovered using the PureLink™ RNA Mini Kit (Invitrogen) according to the manufacturer’s instructions, including on-column DNAse digestion (PureLink™ DNase Set, Invitrogen). RNA quality was assessed using a Bioanalyzer (2100 Bioanalyzer, Agilent). RNA abundance was quantified based on the height of the 28S ribosomal peak since co-purifying contaminants confounded RNA abundance determinations based on total RNA level and RIN.

Twenty-five ng of total RNA was used to construct an RNA-seq library for each sample. RNA was converted to cDNA using the SMART-Seq v4 Ultra Low Input RNA kit. The cDNA was fragmented using a sonication device (Covaris, E200). Library construction was completed using the ThruPLEX® DNA-seq Kit (Rubicon Genomics). Size selection was performed with AMPure XP Beads (Beckmann Coulter) as per the manufacturer’s directions. Quality and mean fragment size of library samples were assessed with the Bioanalyzer prior to sequencing. Libraries were sequenced on a NextSeq 550 deep sequencing unit (150 cycles, paired-end, Illumina) at the Biopolymers Facility at Harvard Medical School, Boston, MA. Reads were mapped using STAR aligner to the *Mus musculus* genome (mm10, Ensembl release 89) [35]. Sequencing and alignment quality was analyzed with FastQC [36], RSeQC [37], and Picard (broadinstitute.github.io/picard). Read counts were calculated using the Subread package.

Because blood, bone marrow, and muscle could not be completely removed from the tibial cortical bone prior to RNA extraction, we computationally removed reads representing 894 transcripts which we had previously shown to comprise ~ 10% of all cortical bone mRNA and come from these contaminating tissues [34]. We then used EdgeR [38] to measure transcript abundance by calculating fragments per kilobase of transcript per million fragments mapped (FPKM).

We quantified fold-changes in expression between *Bglap/2*^dko/dko^ and wild-type transcripts using the DESeq2 package [39]. Reported in Table 2 are those transcripts having a mean FPKM greater than 3 in wild-type, *Bglap/2*^dko/dko^, or in both mice, that change significantly (adjusted p values < 0.05) after correcting for multiple hypothesis testing using the Benjamini-Hochberg method. Supplemental Table 4 includes adjusted p values for all transcripts regardless of FPKM.

## Supporting Information Legends

<u>Supplemental Table 1. Biomechanical Testing on Male *Bglap/2*^dko/dko^ and age-matched wild-type animals</u>. Individual measurements for each mouse are shown for Ultimate Force, Stiffness, and Energy to FU. *Bglap/2*^dko/dko^ (KO/KO, n=12) and wild-type (WT, n=12) are noted and the average and standard deviation are presented for each parameter.

<u>Supplemental Table 2. Biomechanical Testing on Female *Bglap/2*^dko/dko^ and age-matched wild-type animals</u>. Individual measurements for each mouse are shown for Ultimate Force, Stiffness, and Energy to FU. *Bglap/2*^dko/dko^ (KO/KO, n=14) and wild-type (WT, n=18) are noted and the average and standard deviation are presented for each parameter.

<u>Supplemental Table 3. FTIR Imaging Quantitation for Cortical and Trabecular Bone</u>. Quantitation of various parameters for *Bglap/2*^dko/dko^ (KO/KO, n=4) and wild-type (n=3) female mice are noted

<u>Supplemental Table 4. Differential expression calculated using the DESeq2 package of cortical bone mRNA transcripts between male *Bglap/2^dko/dko^* and male wild-type littermate mice</u>. This table includes adjusted p values (controlling for multiple hypothesis testing) for all transcripts regardless of FPKM.

## Acknowledgements

We thank other members of the Williams, Warman, Robling, and Dowd Laboratories for suggestions and technical assistance, and Dr. Nicholas Stylopoulos for advice regarding serial glucose measurements. Key members of the VARI Vivarium and Transgenics Core include Bryn Eagleson, Adam Rapp, Nicholas Getz, Audra Guikema, Tristan Kempston, Malista Powers, and Tina Meringa. This work was funded by the Van Andel Research Institute and NIH grants to BOW (AR068668), MLW and AGR (AR053237), MLW (AR064231), and by a HHMI Medical Research Fellowship to JCWH.

